# A cluster-aware, weighted ensemble clustering method for cell-type detection

**DOI:** 10.1101/246439

**Authors:** Daphne Tsoucas, Guo-Cheng Yuan

## Abstract

Single-cell analysis is a powerful tool for dissecting the cellular composition within a tissue or organ. However, it remains difficult to detect rare and common cell types at the same time. Here we present a new computational method, called GiniClust2, to overcome this challenge. GiniClust2 combines the strengths of two complementary approaches, using the Gini index and Fano factor, respectively, through a cluster-aware, weighted ensemble clustering technique. GiniClust2 successfully identifies both common and rare cell types in diverse datasets, outperforming existing methods. GiniClust2 is scalable to very large datasets.

## INTRODUCTION

Genome-wide transcriptomic profiling has served as a paradigm for the systematic characterization of molecular signatures associated with biological functions and disease-related alterations, but traditionally this could only be done using bulk samples that often contain significant cellular heterogeneity. The recent development of single-cell technologies has enabled biologists to dissect cellular heterogeneity within a cell population. Such efforts have led to an increased understanding of cell-type composition, lineage relationships, and mechanisms underlying cell-fate transitions. As the throughput of single-cell technology increases dramatically, it has become feasible not only to characterize major cell types, but also to detect cells that are present at low frequencies, including those that are known to play an important role in development and disease, such as stem and progenitor cells, cancer-initiating cells, and drug-resistant cells [1, 2].

On the other hand, it remains a computational challenge to fully dissect the cellular heterogeneity within a large cell population. Despite the intensive effort in method development [3–8], significant limitations remain. Most methods are effective only for detecting common cell populations, but are not sensitive enough to detect rare cells. On the other hand, a number of methods have been developed to specifically detect rare cells [9–12], but the features used in these methods are distinct from those distinguishing major populations. Existing methods cannot satisfactorily detect both large and rare cell populations. A naïve approach combining features that are either associated with common or rare cell populations fails to characterize either type correctly, as a mixed feature space will dilute both common and rare cell-type-specific biological signals, an unsatisfactory compromise.

To overcome this challenge, we have developed a new method, GiniClust2, to integrate information from complementary clustering methods using a novel ensemble approach. Instead of averaging results from individual clustering methods, as is traditionally done, GiniClust2 selectively weighs the outcomes of each model to maximize the methods’ respective strengths. We show that this cluster-aware weighted ensemble approach can accurately identify both common and rare cell types and is scalable to large datasets.

## RESULTS

### Overview of the GiniClust2 method

An overview of the GiniClust2 pipeline is shown in Fig. 1. We begin by independently running both a rare cell-type detection method and a common cell-type detection method on the same data set (Fig. 1a). In a previous study [11], we showed that different strategies are optimal for identifying genes associated with rare cell types than for common ones. Whereas the Fano factor is a valuable metric for capturing differentially expressed genes specific to common cell types, the Gini index is much more effective for identifying genes that are associated with rare cells [11]. Therefore, we were motivated to develop a new method that combines the strengths of these two approaches. To facilitate a concrete discussion, here we choose GiniClust as the Gini-index based method and k-means as the Fano-factor based method. However, the same approach can be used to combine any other clustering methods with similar properties. We call this new method GiniClust2.

**Figure 1.**
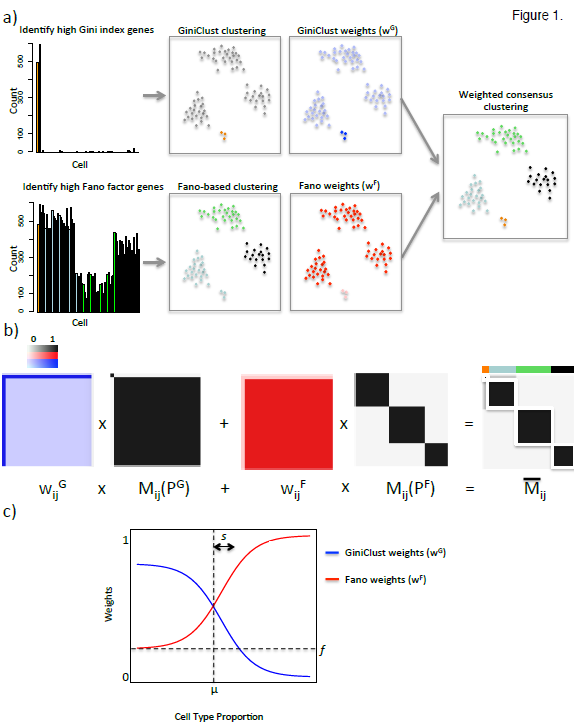
An overview of the GiniClust2 pipeline. (a) The Gini index and Fano factor are used (left), respectively, to select genes for GiniClust and Fano-based clustering (middle left). A cluster-aware, weighted ensemble method is applied to each of these, where cluster-aware weights *w_i_* are represented by the shading of the cells (middle right), to reach a consensus clustering (right). (b) A schematic of the weighted consensus association calculation, with association matrices in black and white, weighting schemes in red and blue, and final GiniClust2 clusters highlighted in white. (c) GiniClust and Fano-based weights, as a function of cell type proportion.

Our goal is to consolidate these two differing clustering results into one consensus grouping. The output from each initial clustering method can be represented as a binary-valued connectivity matrix, M_ij_, where a value of one indicates cells i and j belong to the same cluster (Fig. 1b). Given each method’s distinct feature space, we find that GiniClust and k-means tend to emphasize the accurate clustering of rare and common cell types, respectively, at the expense of their complements. To optimally combine these methods, a consensus matrix is calculated as a cluster-aware, weighted sum of the connectivity matrices (Fig. 1b). Higher weights are assigned to rare clusters over common clusters for GiniClust, whereas higher weights are assigned to common clusters over rare clusters for k-means (Fig. 1c). For simplicity, the weighting functions are modeled as logistic functions which can be specified by three tunable parameters: *μ* is the cluster size at which GiniClust and Fano-factor-based clustering methods have the same detection precision, *s* represents how quickly GiniClust loses its ability to detect rare cell types, and *f* represents the importance of the Fano cluster membership in determining the larger context of the membership of each cell. The values of these parameters are estimated from the data (**Supplemental Information**). An additional round of clustering is then applied to the consensus matrix to identify both common and rare cell clusters. The mathematical details are described in the **Materials and Methods** section.

### Accurate detection of both common and rare cell types in a simulated dataset

We started by evaluating the performance of GiniClust2 using a simulated scRNA-seq dataset, which contains two common clusters (of 2000 and 1000 cells, respectively) and four rare clusters (of 10, 6, 4, and 3 cells, respectively) (**Materials and Methods**, Fig. 2a). We first applied GiniClust and k-means independently to cluster the cells. As expected, GiniClust correctly identifies all 4 rare cell clusters, but merges the two common clusters into a single large cluster (Fig. 2b). In contrast, k-means (with k=2) accurately separates the two common clusters, while lumping together all four rare cell clusters into the largest group (Fig. 2b). Increasing k does not improve the ability to detect rare cells, but rather divides each common cluster into smaller clusters, indicating an intrinsic limitation of selecting gene features using the Fano factor (Supplemental Fig. 1a).

**Figure 2.**
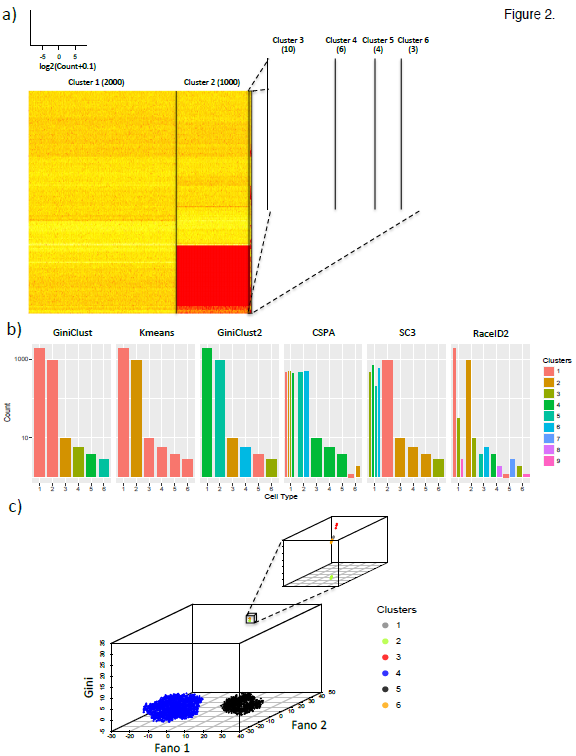
The application of GiniClust2 and competing methods to simulated data. (a) A heatmap representation of the simulated data with 6 distinct clusters, showing the genes permuted to define each cluster. A zoomed-in view of the rare clusters is shown in the smaller heatmap. (b) A comparison between the true clusters (x-axis) and clustering results from GiniClust2 and comparable methods (y-axis). Each cluster is represented by a distinct color bar. Multiple bars are shown if a true cluster is split into multiple clusters by a clustering method. (c) A three-dimensional visualization of the GiniClust2 clustering results using a composite tSNE plot, combining two Fano-based tSNE dimensions and one Gini-based tSNE dimension. Insets show zoomed-in views of the corresponding regions.

We next used the GiniClust2 weighted ensemble step to combine the results from GiniClust and k-means. Of note, all six cell clusters are perfectly recapitulated by GiniClust2 (Fig. 2b), suggesting that GiniClust2 is indeed effective for detecting both common and rare cell clusters. To aid visualization, we created a composite tSNE plot, projecting the cells into a three-dimensional space based on a combination of a two-dimensional Fano-based tSNE map and a one-dimensional Gini-based tSNE map (Fig. 2c). A three-dimensional space is required because, although the Fano-based dimensions are able to clearly separate the two common clusters, the rare clusters are overlapping and cannot be fully discerned. The third (Gini) dimension results in complete separation of the rare clusters.

Since the number of common clusters is unknown in advance, we also tested the robustness of GiniClust2 with respect to other choices of k. We found that setting k=3 provides the same final clustering, while further increase results in poorer performance by splitting of the larger clusters (Supplemental Fig. 1b). By default, the value of k was chosen using the gap statistic [13]. However, this may not be optimal in various cases [14], therefore additional exploration is often needed to select the optimal value for k.

For comparison, we evaluated the performance of two unweighted ensemble clustering methods. First, we used the cluster-based similarity partitioning algorithm (CSPA) [15] to combine the GiniClust and k-means (k=2) clustering results. The consensus clustering splits the common clusters into six subgroups, whereas cells in the four rare clusters are assigned to one of two clusters shared with the largest common cell group (Fig. 2b). Without guidance, the consensus clustering treats all clustering results equally and attempts to resolve any inconsistency via suboptimal compromise. The second method we considered, known as SC3 [4], is specifically designed for single-cell analysis. This method performs an unweighted ensemble of k-means clusterings for various parameter choices without specifically targeting rare cell detection. Regardless of the specific parameter choices, k-means cannot resolve the rarest clusters, and the final ensemble clustering splits the largest group into three and differentiates only one of the four rare clusters (Fig. 2b). These analyses suggest that our cluster-aware, weighted ensemble approach is important for optimally combining the strengths of different methods.

We also compared the performance of GiniClust2 with other rare cell type detection methods. In particular, we compared with RaceID2 [10], which is an improved version of RaceID [9] developed by the same group. For fair comparison, we considered k=2, the exact number of common cell clusters. RaceID2 over-estimated the number of clusters, and split both common and rare cells clusters into smaller subclusters (Fig. 2b). Using other choices of k led to even poorer outcomes. This tendency of over-clustering was consistent with our previous observations [11].

### Robust identification of rare cell types over a wide range of proportions

In order to evaluate the performance of GiniClust2 on analyzing real scRNA-seq datasets, we focused on one of the largest public scRNA-seq datasets generated by 10X Genomics [16]. The dataset consists of transcriptomic profiles of about 68,000 peripheral blood mononuclear cells (PBMCs) [16], which were classified into 11 subpopulations based on transcriptomic similarity with purified cell-types. It was noted that the transcriptomic profiles of several subpopulations are nearly indistinguishable [16].

To reduce the effects of stochastic variation and technical artifacts, we started by considering only a subset of cell types that are distinct from one another. In particular, we focused on three large subpopulations: CD56+ natural killer (NK) cells, CD14+ Monocytes, and CD19+ B cells (Fig. 3a). To systematically compare the ability of different methods in detecting both common and rare cell types, we created a total of 120 random subsamples that mix different cell types at various proportions (Supplementary Table 1), with the rare cell type (NK cells) proportions ranging from 0.4% to 11.6% (see **Materials and Methods** for details).

**Figure 3.**
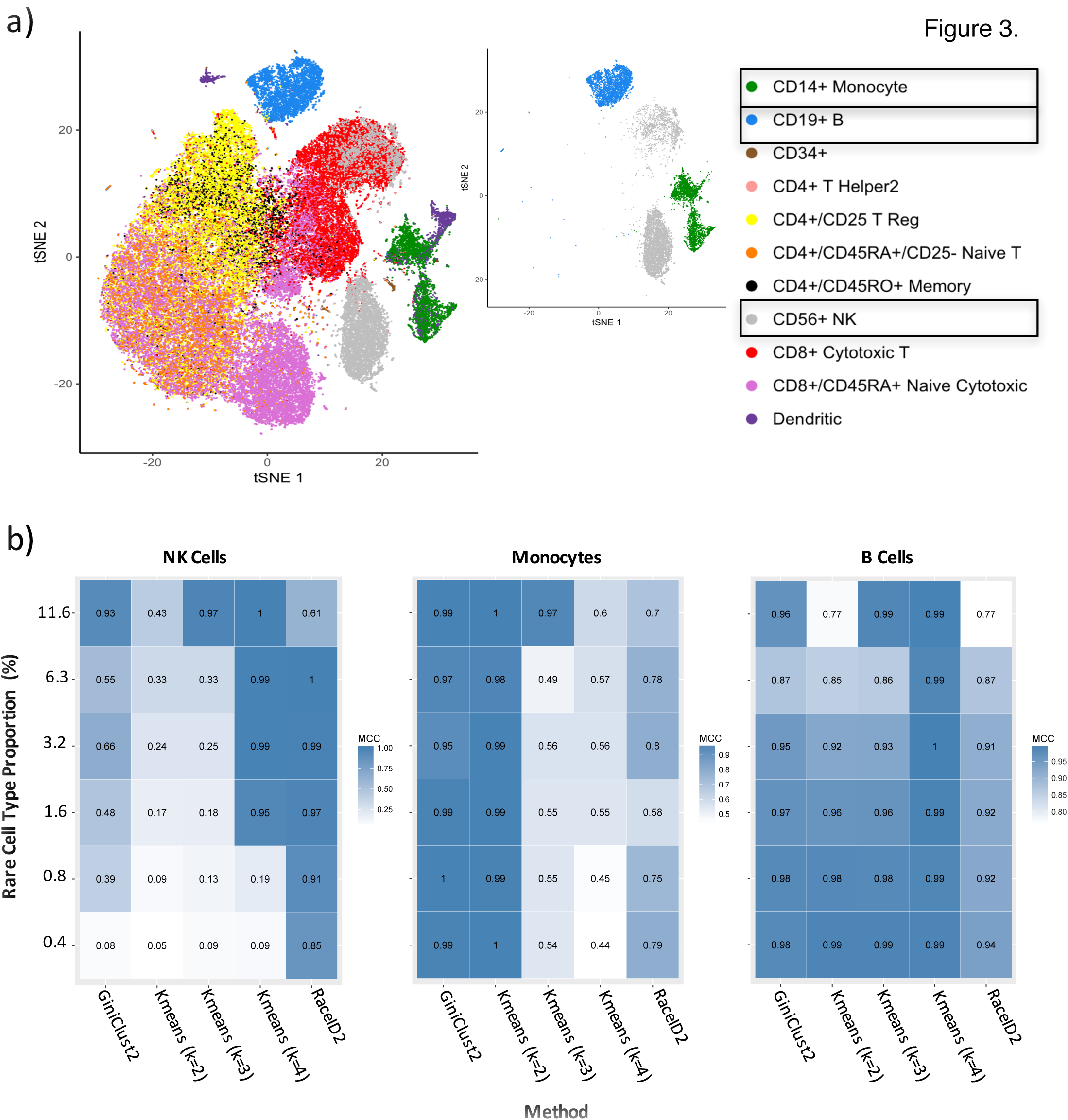
Analysis of the 68k PBMC dataset [16]. (a) A visualization of reference labels for the full data set (left), along with the 3 cell subtypes selected for detailed analysis (right). (b) Comparison of the performance of different clustering methods, quantified by a Matthews correlation coefficient (MCC) [17] for each of the three cell types.

We applied GiniClust2 and comparable methods to the down-sampled datasets generated above. Each method was evaluated based on its ability to detect each cell type using three Matthews correlation coefficients (MCC) [17] (Fig. 3b). The MCC is a metric that quantifies the overall agreement between two binary classifications, taking into account both true and false positives and negatives. The MCC value ranges from −1 to 1, where 1 means a perfect agreement between a clustering and the reference, 0 means the clustering is as good as a random guess, and −1 means a total disagreement between a clustering and the reference.

RaceID2 is the best method for detecting the rare NK cell type at a frequency of 1.6% or lower, and GiniClust2 is the next best method. While the performance of GiniClust degrades as the “rare” cell type becomes more abundant, k-means becomes more powerful in such cases, and results in the near-optimality of GiniClust2 over a wide range of rare-cell proportions. For the common groups, k-means tends to perform better, but only if the parameter is chosen correctly. For example, k-means with k=4 systematically splits the largest Monocyte group and leads to a relatively low MCC value. The low specificity of RaceID2 leads to false rare cell types within the common groups, and consistently low MCC values. On the other hand, GiniClust2 consistently ranks at or near the top among all methods for these common groups. Taken together, these comparative results suggest that GiniClust2 reaches a good balance for detecting both common and rare clusters.

### Detection of rare cell types in differentiating mouse embryonic stem cells

To test if GiniClust2 is useful for detecting previously unknown, biologically relevant cell types, we analyzed a published dataset associated with leukemia inhibitory factor (LIF) withdrawal induced mouse embryonic stem cell (mESC) differentiation [18]. Previously, we applied GiniClust to analyze a subset containing undifferentiated mESCs, and identified a rare group of Zscan4-enriched cells [11]. As expected, these rare cells were rediscovered using GiniClust2.

In this study, we focused on the cells assayed on Day 4 post LIF withdrawal, and tested if GiniClust2 might uncover greater cellular heterogeneity than previously recognized. GiniClust2 identified two rare clusters consisting of 5 and 4 cells respectively, corresponding to 1.80% and 1.44% of the entire cell population. The first group contains 25 differentially expressed genes when compared to the rest of the cell population (MAST likelihood ratio test p-value<1e-5, fold change>2), including known primitive endoderm (PrEn) markers such as *Col4a1, Col4a2, Lama1, Lama2*, and *Ctsl*. These genes are also associated with high Gini index values. Overall there is a highly significant overlap between differentially expressed and high Gini genes (Fisher exact test p-value<1e-18). The second group contains 10 differentially expressed genes (MAST likelihood ratio test p-value<1e-5, fold change>2), including maternally imprinted genes *Rhox6, Rhox9*, and *Sct*, all of which are also high Gini genes. Once again there is a significant overlap between differentially expressed and high Gini genes (Fisher exact test p-value<1e-12). Although these clusters were detected in the original publication [18], this was achieved based on a *priori* knowledge of relevant markers. Here, the strength of GiniClust2 is that it can identify these clusters without previous knowledge.

In addition, GiniClust2 identified 2 common clusters. The first group specifically expresses a number of genes related to cell growth and embryonic development, including *Pim2, Tdgf1*, and *Tcf15* (MAST likelihood ratio test p-value<1e-5, fold change>2), indicating it corresponds to undifferentiated stem cells. The second group is strongly associated with a number of genes related to epiblast cells, including *Krt8, Krt18, S100a6, Tagln, Actg1, Anxa2*, and *Flnc* (MAST likelihood ratio test p-value<1e-5, fold change>2), suggesting this group corresponds to an epiblast-like state. Of note, 114 of the 128 genes (Fisher exact test p-value<1e-88) specifically expressed in this group were selected as high Fano-factor genes, confirming the utility of Fano factor in detecting common cell-types. Both populations were discovered in the original publication [18]. The dissimilarity between these cell types is evident in the heatmap (Fig. 4a) and composite tSNE plot (Fig. 4b).

**Figure 4.**
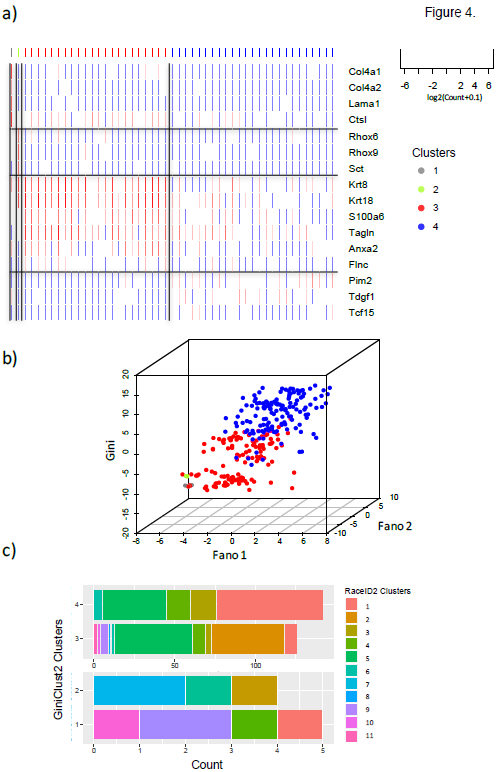
Analysis of the inDrop dataset for day 4 post-LIF mESC differentiation [18]. (a) A heatmap of top differentially expressed genes for each GiniClust2 cluster. The colorbar above the heatmap indicates the cluster assignments. (b) A composite tSNE plot, representing the GiniClust2 results. (c) A comparison of GiniClust2 and RaceID2 clustering results, for common (above) and rare (below) cell types. The same color-coding scheme is used in all panels.

For comparison, we applied RaceID2 to analyze the same dataset. Unlike GiniClust2, RaceID2 broke each cluster into multiple subclusters, and failed to identify the rare cell clusters (Fig. 4c). With k=2, RaceID2 found a total of 11 clusters. Clusters 1, 2, 4, and 9 display an epiblast-like signature, clusters 7, and 10 overexpress genes relating to maternal imprinting, and clusters 8 and 11 correspond to PrEn cells. From these results it appears that RaceID2 has difficulty in differentiating rare, biologically meaningful cell types from outliers.

### Scalability to large data sets

With the rapid development of single-cell technologies, it has become feasible to profile thousands or even millions of transcriptomes at single-cell resolution. Thus, it is desirable to develop scalable computational methods for single-cell data analysis. As a benchmark, we applied GiniClust2 to analyze the entire 68k PBMC data set [16] described above to uncover hidden cell types. The complete analysis took 2.3 hours on a computing cluster with 56 CPUs, 28 cores, and 660 GB memory. For comparison, RaceID2 analysis could not be completed for this large dataset. One possible explanation is this method may be limited to handling data sets with less than 65,536 data points due to an intrinsic restriction in R.

Our analysis identified 9 common clusters and two rare clusters (Fig. 5a), In general, the results of GiniClust2 and k-means are similar; both agree well with the reference cell types (Fig. 5b, NMI = 0.540 for GiniClust2, NMI = 0.553 for k-means). Most of the discrepancy between the clustering results and reference labels are associated with T-cell subtypes. As noted by the original authors [16], these subtypes are difficult to separate because they share similar gene expression patterns and biological functions. The common clusters detected by GiniClust2 and k-means express marker genes known to be specific to the cell types represented in the reference [19] (Fig. 5c).

**Figure 5.**
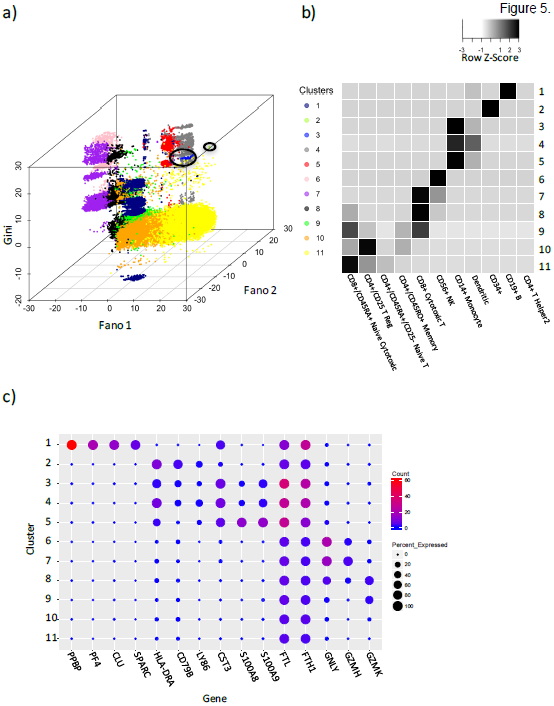
Results from the full 68k PBMC data analysis. (a) A composite tSNE plot of the GiniClust2 results; rare cell types are circled. (b) A confusion map showing similarities between GiniClust2 clusters and reference labels. Values represent the proportion of cells per reference label that are in each cluster. (c) A bubble plot showing expression of cluster-specific genes. Size represents the percentage of cells within each cluster with non-zero expression of each gene, while color represents the average normalized UMI counts for each cluster and gene.

With respect to rare cell types, our first group contains a homogeneous and visually distinct subset of 171 of 262 total CD34+ cells (Cluster 2, Fig. 5a). This cluster was partially detectable using k-means, although it was partially mixed with major clusters. The second rare cell cluster is previously unrecognized (Cluster 3, Fig. 5a). This cluster contains 118 cells (0.17%) within a large set of 5433 immune cells with similar gene expression patterns. Among these 118 cells, 101 cells are classified as monocytes, whereas 16 are classified as dendritic cells, and 1 is classified as a CD34+ cell. Differential expression analysis (MAST likelihood ratio test p-value<1e-5, fold change>2) identified 187 genes that are specifically expressed in this cell cluster, including a number of genes associated with tolerogenic properties, such as *Ftl, Fth1*, and *Cst3* [20], suggesting these cells may be associated with elevated immune response and metabolism. Taken together, these results strongly indicate the utility of GiniClust2 in analyzing large single-cell datasets.

## DISCUSSION AND CONCLUSIONS

According to the “no free lunch” theorems [21], an algorithm that performs well on a certain class of optimization problems is typically associated with degraded performance for other problems. Therefore it is expected that clustering algorithms optimized for detecting common cell clusters are unable to detect rare cell clusters, and vice versa. While ensemble clustering is a promising strategy to combine the strengths of multiple methods [15, 4, 5], our analysis shows that the traditional, unweighted approach does not perform well.

To optimally combine the strengths of different clustering methods, we have developed GiniClust2, which is a cluster-aware, weighted ensemble clustering method. GiniClust2 effectively combines the strengths of Gini-index- and Fano-factor-based clustering methods for detecting rare and common cell clusters, respectively, by assigning higher weights to the more reliable clusters for each method. By analyzing a number of simulated and real scRNA-seq datasets, we find that GiniClust2 consistently performs better than other methods in maintaining the overall balance of detecting both rare and common cell types. This weighted approach is generally applicable to a wide range of problems.

GiniClust2 is currently the only rare-cell-specific detection method equipped to handle such large data sets, as demonstrated by our analysis of the 68k PBMC dataset from 10X Genomics. This property is important for detecting hidden cell types from large datasets, and may be particularly useful for annotating the Human Cell Atlas [22].

## MATERIALS AND METHODS

### Data Source and Preprocessing

Mouse ESC scRNA-seq data was obtained from Gene Expression Omnibus (GEO) with the accession number GSE65525 [18]. This processed data is represented as UMI filtered-mapped counts. Removing genes expressed in fewer than 3 cells, and cells expressing fewer than 2000 genes, we were left with a total of 8055 genes and 278 cells.

The 68k PBMC dataset was obtained from the 10X Genomics website: http://support.10xgenomics.com/single-cell/datasets. The processed data (represented as UMI counts) was filtered and normalized using the code provided by 10X Genomics (https://github.com/10XGenomics/single-cell-3prime-paper). The resulting data consists of a total of 20387 genes and 68579 cells. Cell-type labels were assigned based on the maximum correlation between the gene expression profile of each single cell to 11 purified cell populations, using the code provided by 10X Genomics.

### GiniClust2 method details

GiniClust2 has been implemented in R and deposited at Github with URL https://github.com/dtsoucas/GiniClust2. The GiniClust2 pipeline contains the following steps.

#### Step 1: Clustering cells using Gini-index based features

The Gini index for each gene is calculated and normalized as described before [11]. Briefly, the raw Gini index is calculated as twice the area between the diagonal and the Lorenz curve, taking a range of values between 0 and 1. Raw Gini index values are normalized by removing the trend with maximum expression levels using a two-step LOESS regression procedure as described in [11]. Genes whose normalized Gini index is significantly above zero (p-value < 0.0001) are labeled high Gini genes and selected for further analysis.

A high-Gini-gene-based distance is calculated between each pair of cells using the Jaccard distance metric. This is used as input into DBSCAN [23], which is implemented using the dbscan function in the fpc R package, with method= “dist”. Parameter choices for eps and MinPts are discussed in the **Supplemental Information**.

#### Step 2: Clustering cells using Fano-factor based features

The Fano factor is defined as the variance over mean expression value for each gene. The top 1000 genes are chosen for further analysis. Principal component analysis (PCA) is applied to the gene expression matrix for dimensionality reduction, using the svd function in R. The first 50 principal components are reserved for clustering analysis. Cell clusters are identified by k-means clustering, using the kmeans function in R with default parameters. Optimal choice of k is discussed in the **Supplemental Information**. To improve robustness, 20 independent runs of k-means clustering with different random initializations are applied to each dataset, and the optimal clustering result is selected.

#### Step 3. Combining the results from Steps 1 and 2 via a cluster-aware, weighted ensemble approach

We adapted the weighted consensus clustering algorithm developed by Li and Ding [24] by further considering cluster-specific weighting. For GiniClust, higher weights are assigned to the rare cell clusters and lower weights to common clusters, whereas the opposite scheme is used to weight the outcome from k-means clustering. This allows us to combine the strength of each clustering method. The mathematical details are described as follows, and visualized in Fig. 1b.

Let *P^G^* be the partitioning provided by GiniClust, and *P^F^* the partitioning provided by Fano-factor-based clustering. Each partition consists of a set of clusters: 
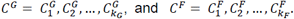
 Define the connectivity matrices as:

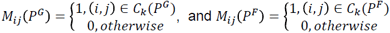

If two cells are clustered together in the same group, their connectivity is 1, while if they are clustered separately, their connectivity is 0. Define the weighted consensus association as:

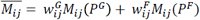

where 
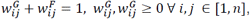
 *n* represents the number of cells. Weights 
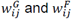
 are specific to each pair of cells, and are determined based on 
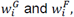
 on weights that are specific to each cell. These cell-specific weights are determined based on the rarity of the cell cluster that the cell is assigned to, the specifics in which are discussed in the next section. The cell-pair-specific weights are translated from the cell-specific weights as follows:

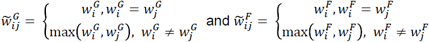

Then, weights are normalized to 1:

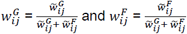

Each cell-cell pair will thus be assigned a weighted consensus association between 0 and 1, which is a cell-type-proportion-based weighted average of both GiniClust and Fano-factor-based clustering associations.

At this point, the weighted consensus association matrix provides a probabilistic clustering for each cell, where each entry represents the probability that cell i and cell j reside in the same cluster. To transform this into a final deterministic clustering assignment, we optimize the following:

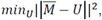

where *U* is any possible connectivity matrix. In Li and Ding [24], this optimization problem is solved via symmetric non-negative matrix factorization (NMF) to yield a soft clustering. To obtain a hard clustering we add an orthogonality constraint, leading to orthogonal NMF, or k-means clustering [25], implemented once again using the kmeans R function.

### Weighting scheme parameters

Cell-specific GiniClust 
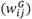
 and Fano-factor-based 
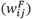
 weights are determined as a function of the size of the cluster containing the particular cell. To illustrate the origins of the weighting scheme, we turn to a conceptual example.

As the proportion of the rare cell type increases, the utility of GiniClust begins to decline. In a complementary manner, the ability of Fano-factor-based k-means clustering to detect the rare cell type is expected to rise. We find this phenomenon to hold true in all the datasets we analyzed, and to be best demonstrated in the subsampled PBMC data results. Supplemental Fig. 4a shows a schematic of the respective abilities of GiniClust and Fano-factor-based k-means clustering to detect a rare cell type over various rare cell type proportions, created by applying smoothing to the subsampled PBMC results. We next calculate the ratio between this GiniClust detection ability and the sum of both GiniClust and k-means detection abilities (Supplemental Fig. 4b). This is a measure of the power of GiniClust over k-means in detecting the rare cell type. When this ratio is 1, GiniClust is solely able to detect the rare cell type, while at zero k-means alone is useful. This curve closely follows a logistic shape, and thus the following equation is used to approximate it:

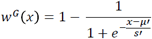

where *μ′* is the rare cell type proportion at which GiniClust and Fano-factor-based clustering methods have the same ability to detect rare cell types, and *s′* represents how quickly GiniClust loses its ability to detect rare cell types above *μ′* (see **Supplemental Information** for more details). Supplemental Fig. 4b also contains this fitted curve, which is then directly used to determine the GiniClust weights 
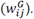
 Fano-factor-based clustering weights are determined by recognizing that while Fano-factor-based clusters may not contain rare cell clusters, they are still indicative of where these rare cells lie in the context of the larger cell population. As such, weights are set such that 
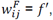
 where *f′* is a constant corresponding to the importance of larger Fano cluster membership for rare cells (see *Supplemental Information* for details on the choice of *f′*). Clustering results tend to be robust to the specific choice of *μ′*, *s′*, and *f′*. Fig. 1c shows the result of subsequent scaling of Gini and Fano weights to sum to one, and their associated scaled parameters *μ, s*, and *f*.

### tSNE visualization

Dimension reduction by tSNE [26] is performed using the Rtsne R package. The tSNE algorithm is first run using the Gini-based distance to obtain a one-dimensional projection of each cell. For large data sets, tSNE is run on the first 50 principal components of the Gini-based distance to prevent tSNE from becoming prohibitively slow. Then, the tSNE algorithm is run using the first 50 principal components of our Fano-based Euclidean distance to obtain a separate two-dimensional projection. The three resulting dimensions (one for Gini-based distance and two for Fano-based distance) are plotted to visualize cluster separation.

### Differential expression analysis on resulting clusters

Differentially expressed genes for each cluster are determined by comparing their gene expression levels to all other clusters. This is performed using the zlm.SingleCellAssay function in the R MAST package [27], with method= “glm”. P-values for differentially expressed genes are calculated using the lrTest function, with a hurdle model.

### SC3 analysis

SC3 [4] was accessed through the SC3 Bioconductor R package. SC3 was applied to the simulated data set post-filtering using default parameters, with k=6 to match the true number of clusters.

### CSPA analysis

Matlab code for the CSPA [15] was accessed through http://strehl.com/soft.html, under “ClusterPack_V2.0.” CSPA was applied to the Gini and Fano-based clustering results for the simulated data set, using the clusterensemble function, specifying the CSPA option. K was specified as 6 to match the true number of clusters.

### RaceID2 analysis

RaceID2 [10] R scripts were accessed through https://github.com/dgrun/StemID. RaceID2 was applied to already-filtered data sets as above to make results directly comparable to GiniClust2, with default parameters. k was set to match the corresponding GiniClust2 k parameter specification.

### Simulation details

We created synthetic data following the same approach as Jiang et al. [11], specifying one large 2000 cell cluster, one large 1000 cell cluster, and four rare clusters of 10, 6, 4 and 3 cells, respectively. Gene expression levels are modeled using a negative binomial distribution, and distribution parameters are estimated using an intestinal scRNA-seq data set using a background noise model as in Grün et al. [9]. To create clusters with distinct gene expression patterns, we permute 100 lowly (mean<10 counts) and 100 highly (mean>10 counts) expressed gene labels for each cluster (see Jiang et al. [11] for more details). This results in a 23,538 gene by 3023 cell data set. After filtering (as above) we are left with 3708 genes and 3023 cells.

### 10X Genomics Data Subsampling

The full 68k 10X Genomics PBMC dataset is down-sampled for model evaluation. We created 6 sets of 20 subsampled data sets each for a total of 120 data sets in the following manner: five cells were randomly sampled from the CD56+ NK cell population to form a “rare” cell group for all 120 datasets. Then, for each set of 20 data sets, cells were randomly sampled from the CD14+ Monocyte and CD19+ B cells in specified numbers to form “common” cell clusters, the details of which are listed in Supplemental Table 1.

## ABBREVIATIONS

tSNE: t-distributed stochastic neighbor embedding
PCA: principal component analysis
CSPA: cluster-based similarity partitioning algorithm
RaceID: rare cell type identification
PBMC: peripheral blood mononuclear cell
NK: natural killer
MCC: Matthews correlation coefficient
LIF: leukemia inhibitory factor
mESC: mouse embryonic stem cell
PrEn: primitive endoderm
NMI: normalized mutual information
MAST: model-based analysis of single-cell transcriptomics
scRNA-seq: single-cell RNA-sequencing
DBSCAN: density-based spatial clustering of applications with noise
NMF: non-negative matrix factorization

## DECLARATIONS

### Ethics approval and consent to participate

Not applicable.

### Consent for publication

Not applicable.

### Availability of data and materials

GiniClust2 has been implemented in R and deposited at Github with URL https://github.com/dtsoucas/GiniClust2. Datasets analyzed were all from publically available sources, and are described in **Methods and Materials** section.

### Authors’ contributions

DT and GCY conceived of and designed the computational method. DT implemented the method. DT and GCY wrote the manuscript. All authors read and approved the final manuscript.

## Acknowledgements

We thank Dr. Lan Jiang and members of the Yuan Lab for helpful discussions, as well as Drs. John Quackenbush and Martin Aryee for their support and advice.

## SUPPLEMENTAL INFORMATION

### Parameter choice for DBSCAN

DBSCAN has two parameters: MinPts and eps. MinPts is specified as 3 for all data sets except for the PBMC data, where MinPts is set to 100, in accordance with the larger size of this data set. This corresponds to the minimum cluster size for which we would expect to see a biologically relevant cluster. In general, we find an appropriate MinPts specification to be about 0.1% of the total number of cells.

The eps parameter is determined by a k-nearest-neighbors (kNN) plot as recommended by the authors of DBSCAN [23]. According to their approach, distance from each point to its kth nearest neighbor is plotted in ascending order, where k=MinPts. This will form a line featuring an inflection point, at which lies the recommended choice for eps [23]. If multiple inflection points exist, this may suggest that multiple values of eps are worth exploring; however, in our case as we are concerned with rare clusters, we only consider the smallest choice of eps, corresponding to the first inflection point. Here, our Gini-based distance metric is particularly low-dimensional due to the use of Jaccard distance and a small number of high Gini genes. This causes cells with similar expression profiles to have pairwise distances of zero, which distorts the traditional kNN-distance curve shape and makes the inflection point harder to visualize (**Supplemental Figure 3**). We provide an alternate numerical approach to approximating the inflection point: after removing all zero distances, the inflection point roughly corresponds to the kNN-distance of the (0.00125*total number of cells*MinPts)^th^ cell. For the 68k data set, the computation of all kNN distances was prohibitive, so we subsampled 2057 cells and computed 3-NN distances to maintain the ratio of MinPts to the total sample size. All analyses of PBMC subsampled data use this same resulting value of eps.

### Choice of k for k-means clustering

We give the option of automatically determining k using the gap statistic. However, observing differentially expressed genes and visualizing k-means clusters gives the best intuition as to the optimal k. We also do not suggest using the gap statistic for large data sets due to its computational demands. For the synthetic data, we chose k=2, in accordance with the number of large clusters, but show that k=3 will also yield the same result (**Supplemental Fig. 1**). For subsampled PBMC data sets, k was chosen as 2 or 3 depending on the ability of k-means to pick up the rare NK cell group. K was chosen as 2 for the day 4 post-LIF mouse embryonic stem cell data as we found this number to best group biologically meaningful cell types.

For the 68k PBMC data, we chose k=10 to allow for direct comparison with clustering results from Zheng et al. [16]. For comparison to the k=10 parameter choice, we additionally show results for both choices k=8 and k=12 for the Fano-based clustering step. All parameter choices perform comparably, with NMIs of 0.542, 0.541 and 0.498, respectively, when compared to the reference labels. The k=8 clustering results in two fewer clusters within the CD56+ NK, CD8+ Cytotoxic T, CD8+/CD45RA+ Naïve Cytotoxic continuum. The k=12 clustering further splits clusters containing overlapping CD8+/CD45RA+ Naïve Cytotoxic, CD4+/CD45RA+/CD25- Naïve T, CD4+/CD25 T Reg, and CD4+/CD45RO+ Memory cells, as well as adding another cluster to the aforementioned CD56+ NK, CD8+ Cytotoxic T, CD8+/CD45RA+ Naïve Cytotoxic continuum. These changes are minor as they occur predominantly in regions of unclear identity. Supplemental Fig. 2 shows the Fano-based tSNE dimension reduction labeled with k-means clusters for k=8, 10 and 12.

### Parameter choices for weighted consensus clustering

Weighting parameters *μ′*, *s′*, and *f′* are determined by the curve:

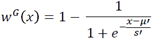

and are represented in **Supplemental Fig. 5**. *μ′* is the rare cell type proportion at which GiniClust and Fano-factor-based clustering methods have the same ability to detect rare cell types. We set this value as 4*(MinPts/total number of cells), where MinPts is the DBSCAN parameter for GiniClust. The smallest possible rare cell type proportion for GiniClust is set as (MinPts/total number of cells). *s′* represents how quickly GiniClust loses its ability to detect rare cell types above *f′*. More quantitatively, we define the 99^th^ percentile of the GiniClust weighting distribution as the point in which GiniClust no longer reliably detects rare cell types. We set *s′* such that the 99^th^ percentile is reached at 6*(MinPts/total number of cells). We expect that at six times the size of smallest possible rare cell type for GiniClust, any cell type found likely consists of a heterogeneous grouping of cells. *f′* represents the importance of the Fano cluster membership in determining the larger context of the membership of each cell, and by default *f′* = 0.1. We find that final membership is not sensitive to small changes in *f′*.

**Supplemental Table 1.**
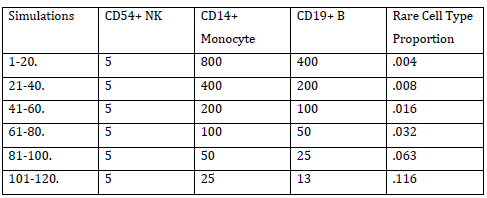
Cell numbers in three different cell types for each of 120 subsampled datasets from 68k PBMCs.

## FIGURE LEGENDS

**Supplemental Figure 1**. The effect of various choices of k on (a) the k-means step and (b) the overall clustering of our GiniClust2 method for the simulated data. Each bar represents the contribution of a cluster to the total number of cells in each reference type.

**Supplemental Figure 2**. The effect of various choices of k on the k-means step for the full 68k PBMC data.

**Supplemental Figure 3**. An illustration of the eps selection process for DBSCAN. Eps is chosen as the distance at the inflection point in the k-nearest-neighbors distance plot.

**Supplemental Figure 4**. Weighting scheme details. (a) Rare cell type detection abilities of GiniClust and k-means over a range of rare cell type proportions. (b) A representation of the ability of GiniClust to detect rare cell types over k-means, and its logistic fit. Parameters *μ′* and *s′* determine the shape of the curve.

